# Genetic diversity and evolution of Hantaan virus in China and its neighbors

**DOI:** 10.1101/2020.01.28.922724

**Authors:** Naizhe Li, Aqian Li, Yang Liu, Wei Wu, Chuan Li, Dongyang Yu, Yu Zhu, Jiandong Li, Dexin Li, Shiwen Wang, Mifang Liang

**Affiliations:** Key Laboratory of Medical Virology and Viral Diseases, Ministry of Health of People’s Republic of China, National Institute for Viral Disease Control and Prevention, Chinese Center for Disease Control and Prevention, Beijing, 102206, China; Department of Microbiology, Anhui Medical University, Hefei, 230032, China; Center for Biosafety Mega-Science, Chinese Academy of Sciences, Beijing 100049, P. R. China

**Keywords:** Hantaan virus, recombination, Bayesian phylogenetics, Viral phylodynamic, Spatial transmission

## Abstract

**Backgroud:** Hantaan virus *(HTNV)*, as one of the pathogenic hantaviruses of HFRS, has raised serious concerns in Eurasia. China and its neighbors, especially Russia and South Korea, are seriously suffered HTNV infections. Recent studies reported genetic diversity and phylogenetic features of HTNV in different parts of China, but the analyses from the holistic perspective are rare.

**Methodology and Principal Findings:** To better understand HTNV genetic diversity and dynamics, we analyzed all available complete sequences derived from the S and M segments with bio-informatic tools. Our study revealed 11 phylogroups and sequences showed obvious geographic clustering. We found 42 significant amino acid variants sites and 18 of them located in immune epitopes. Nine recombination events and seven reassortment isolates were deteced in our study. Sequences from Guizhou were highly genetic divergent, characterized by the emergence of multiple lineages, recombination and reassortment events. We found that HTNV probably emerged in Zhejiang about 1,000 years ago and the population size expanded from 1980s to 1990s. Bayesian stochastic search variable selection analysis revealed that Heilongjiang, Shaanxi and Guizhou played important roles in HTNV evolution and migration.

**Conclusions/Significance:** These findings reveal the original and evolution features of HTNV which might assist in understanding Hantavirus epidemics and would be useful for disease prevention and control.

**Author summary:** Hemorrhagic Fever with Renal Syndrome (HFRS) and Hantavirus Pulmonary Syndrome (HPS) are endemic zoonotic infectious diseases caused by hantaviruses that belong to the Family Bunyaviridae. Hantaviruses have gained worldwide attention as etiological agents of emerging zoonotic diseases, with fatality rates ranging from <10% up to 60%. However, our knowledge about the emergence and evolution of HTNV is limited. To get more information about HTNV genetic diversity and phylogenetic features in holistic perspective, we investigated the genetic diversity and spatial distribution of HTNV using all available whole genomic sequences of S and M segments. We also gain insights into the genetic diversity and spatial-temporal dynamics of HTNV. These data can augment traditional approach to infectious disease surveillance and control.

## Introduction

Hantaviruses have gained worldwide attention as etiological agents of emerging zoonotic diseases, namely hemorrhagic fever with renal syndrome (HFRS) in Eurasia and hantavirus cardiopulmonary syndrome (HCPS) in Americas, with fatality rates ranging from <10% up to 60%[1]. Among all the countries, China is the most seriously affected one which account for over 90% of the total HFRS cases around the world[2, 3]. It has been reported that the HFRS death rate in China was 2.89% during the years 1950– 2014[4].

Although a declining HFRS trend has been observed at a global scale in China, there still exist certain local regions that continue to display increasing HFRS trends [5]. However, the causative agent of the disease remained unknown until the early 1970s, when Lee et al. reported on Hantaan virus (HTNV), present in the lungs of its natural reservoir, the striped field mouse (Apodemus agrarius)[6].

Hantaan virus, as one of the pathogenic hantaviruses of HFRS, is a member of genus Orthohantavirus, family Hantaviridae in the order Bunyavirales. The virus genome consists of three separate segments of negative-stranded RNA referred to as small (S), medium (M), and large (L) segments, which encode nucleocapsid protein (NP), two envelope glycoproteins (Gn and Gc) and viral RNA-dependent RNA polymerase (RdRP), respectively[7]. Recent studies[8–10] reported genetic diversity and phylogenetic features of HTNV in different parts of China, but the analyses from the holistic perspective are rare. Also, the mechanisms underlying the emergence and evolution of HTNV are poorly understood. Understanding the phylogenetic factors contributing to HTNV transmission has important implications for our understanding of its epidemiology and providing insights about the surveillance and outbreak predictions or preparation.

In this study, we focused on the genetic diversity and evolution history of Hantaan virus. Whole-genome sequences were analyzed with bio-information tools. The results revealed the phylogenetic relationships among HTNV strains with both Bayesian and Maximum Likelihood methods. To deduce geographic origins and migration patterns of HTNV epidemics, BEAST software was used in this study. Furthermore, the recombination and reassortment events were also detected.

## Materials and methods

### 1. Sequence dataset

All the complete S gene and M gene sequences of Hantaan orthohantavirus deposited until June 2019 were collected from Virus Pathogen Database and Analysis Resource (ViPR, www.viprbrc.org).

Only one sequence was retained for the identity sequences with the same strain names. It should be mentioned that sequences from Shandong province (accession no. KY639536-KY639711) were not defined as HTNV by ViPR and Genebank but can be organized in Hantaan orthohantavirus (see below). These sequences were still analyzed in this study. All the sequences were aligned using MAFFT version7[11] with default settings followed by manual refinement. The coding sequences were retained and used for the following analyses[12].

### 2. Recombination detection

As recombination seriously affects phylogenetic inference, the whole dataset was tested for the presence of recombination signals using RDP4[13]. RDP, GENECONV, Maximum Chi-squared, Chimaera, Bootscan, Sister Scanning and 3Seq were used Only events with P values<0.01 that were confirmed by four or more methods were considered as recombination.

### 3. Phylogenetic analyses and amino acid substitution analyses

To reduce the potential bias, the recombination segments were removed from the datasets firstly. The phylogenetic relationships of the complete S and M gene sequences were estimated using a Bayesian Markov Chain Monte Carlo (MCMC) method as implemented in MrBayes v3.2.2[14] and a maximum-likelihood (ML) phylogenetic inference as implemented in IQ-TREE v1.6.8[15]. Dobrava virus (DOBV), Puumala virus (PUUV), Sin Nombre virus (SNV) and Thottapalayam virus (TPMV) were designated as outgroup.

For Bayesian MCMC analysis, the suitability of substitution models for our datasets were assessed using jModelTest v2.1.10[16], which performed a statistical model selection procedure based on the Akaike Information Criterion (AIC). It identified the best fitting substitution model GTR+Γ for both datasets of complete S and M gene sequences. The

MCMC settings consisted of two simultaneous independent runs with four chains each, which were run for 20 million generations and sampled every 200 generation with a 25% burn-in.

For maximum-likelihood analysis, the best-fitting nucleotide substitution model that minimizes the Bayesian Information Criterion (BIC) score was selected by ModelFinder[17], implemented in IQ-TREE. A ML tree was constructed using the best-fitting nucleotide the substitution model, and statistical robustness of the branching order within the tree was assessed using ultrafast bootstrap support values[18] (1000 replicates) and the SH-like approximate likelihood ratio test[19] (SH-aLRT) (5000 replicates). The trees were visualized in FigTree v1.4.4 (http://tree.bio.ed.ac.uk/software/figtree/).

Metadata-driven comparative analysis tool (meta-CATS) supplied by ViPR was used to analyze the variation of amino acid between the different phylogenetic groups, host and sample collection years. The P value threshold was set to 0.05. If a specific amino acid exists only in one species or one group but not in other species or group, this amino acid is considered a ‘significant amino acid’ marker (or synapomorphy).

### 4. Coalescent and evolution analyses

Complete S gene sequences were used to deduce the evolution history of Hantaan virus. To avoid potential biases due to sampling heterogeneity, the dataset was reduced using CD-HIT[20] by clustering together sequences with a nucleotide sequence identity threshold of 99%. Temporal evolutionary signal in ML tree was evaluated by using TempEst[21] v1.5.1, which plots sample collection dates against root-to-tip genetic distances obtained from the ML phylogeny tree.

The tMRCA, substitution rates, and evolution history were estimated using a Bayesian serial coalescent approach implemented in BEAST v1.10.4[22]. jModelTest v2.1.10. was used to determine the model of nucleotide substitution that best fit the dataset, and the dataset was subsequently run using GTR+I+Γ. Both strict and relaxed (uncorrelated exponential and uncorrelated lognormal) molecular clocks were used for the analysis combined with different tree prior (Constant Population size, GMRF Bayesian Skyride, Bayesian Skyline were used in this study). Time-scale was inferred using an informative substitution rate interval (1.0х10^4^ to 1.0х10^3^ substitutions/site/year) previously estimated for the N gene of rodent-borne hantavirus[23]. Models were compared pairwise by estimating the Log marginal likelihood via path sampling (PS) and stepping stone (SS) analysis[24–26]. For each model, an MCMC was run for 100 million generations, sampling every 10,000 steps. Posterior probabilities were calculated using the program Tracer1.7 after 10% burn-in. The results were accepted only if the effective sample sizes (ESS) exceeded 200.

The selected molecular clock/demographic model (strict clock with Bayesian skyline prior in this study) was then used for the Bayesian phylogeographic inference based on the continuous-time Markov Chain (CTMC) process over discrete sampling locations to estimate the diffusion rates among locations, with Bayesian stochastic search variable selection (BSSVS). The MCMC was run for 200 million generations, with sampling every 20,000 generations. Only parameter estimates with ESS values exceeding 200 were accepted. A maximum clade credibility (MCC) tree with the phylogeographic reconstruction was selected from the posterior tree distribution after a 10% burn-in using the Tree Annotator v 1.10 and was manipulated in FigTree v1.4.4. The migration routes were visualized using SPREAD3[27]. Migration pathways were considered to be important when they yielded a Bayes factor greater than 15 and when the mean posterior value of the corresponding was greater than 0.50. Bayes factors were interpreted according to the guidelines of Kass and Raftery[28].

## Results

### 1. HTNV sequences dataset

A total of 225 complete S gene sequences and 180 complete M gene sequences from 238 HTNV strains were contained in the dataset. The sampling dates of the sequences ranged from 1976 to 2017. According to the sample collection area, 18 regions were defined, including Russia, South Korea and 16 provinces of China (Fig 1A). The samples were collected from 9 kinds of hosts, including *Apodemus agrarius, Apodemus peninsulae, Rattus norvegicus, Mus musculus, Nivienter confucianus, Tscherskia triton, Mirotus fortis, Crocidura russula, and Homo sapiens.* The details of the sequences are shown in Fig 1.

**Fig 1.**
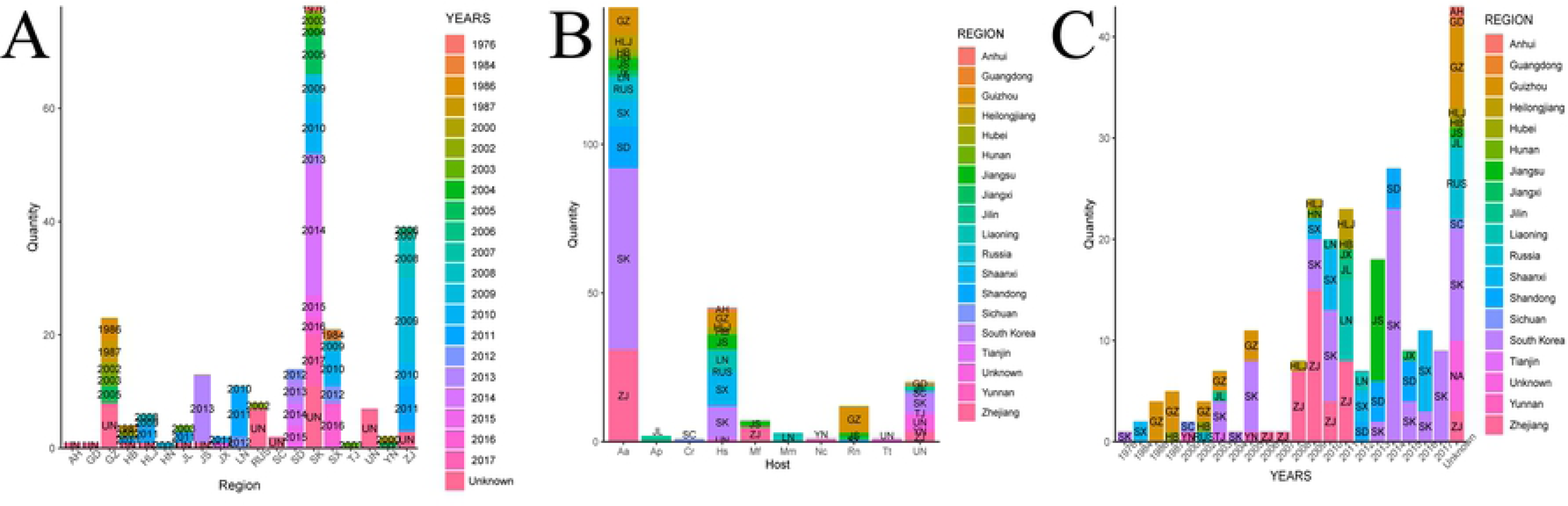
The details of the datasets. (A) The number of isolates collected from different regions. The bars were colored in different sampling years. (AH, Anhui; GD, Guangdong; GZ, Guizhou; HB, Hubei; HLJ, Heilongjiang; HN, Hunan; JL, Jilin; JS, Jiangsu; JX, Jiangxi; LN, Liaoning; RUS, Russia; SC, Sichuan; SD, Shandong; SK, South Korea; SX, Shaanxi; TJ, Tianjin; UN, Unknown; YN, Yunnan; ZJ, Zhejiang;) (B) The number of isolates collected from different hosts. (Aa, Apodemus agrarius; Ap, Apodemus peninsulae; Rn, Rattus norvegicus; Mm,Mus musculus; Nc, Nivienter confucianus; Tt, Tscherskia triton; Mf, Mirotus fortis; Cr, Crocidura russula; Hs, Homo sapiens.) The bars were colored in different regions. (C) The number of isolates collected in different years.

### 2. Recombination events detection

To reduce the potential bias, the recombination segments should be removed from the datasets. Recombination events were detected among all the 225 complete S gene sequences and 180 complete M gene sequences from 238 HTNV strains collected in this study. The RDP analysis suggested 9 recombination events in 6 HTNV isolates, and all the recombination events were occurred in M gene segment (Table 1). Similar recombination events were detected in three isolates (CGAa31MP7, CGAa31P9, CGHu2). All this three isolates were collected during 2004 to 2005 in Guizhou province[8]. It would imply that these isolates have the same ancestor which has experienced recombination before. Although only 5 methods suggested the recombination event occurred in strain JN131026, the p-values were all lower than 0.01. Considering the consensus score is higher than 0.6 (0.646), we defined it as a recombinant isolate. P-values of the event tested in strain A16 were in high credible level.

**Table 1.** Recombination events detected in all HTNV sequences

| Events | Recombinant | Segment | Beginning | Ending | Major Parent | Minor Parent | Method(s) by which breakpoint was detected in RDP4 |
| --- | --- | --- | --- | --- | --- | --- | --- |
| 1 | CGAa31MP7 | M | 1756 | 1916 | CGRni1 | CGRn15 | RDP/GENECONV/MaxChi/Chimaera/Bootscan/SisScan/3Seq |
| 2 | CGAa31MP7 | M | 2315 | 3402 | Q32 | CGRn2616 | RDP/GENECONV/MaxChi/Chimaera/Bootscan/SisScan/3Seq |
| 3 | CGAa31P9 | M | 1756 | 1916 | CGRni1 | CGRn15 | RDP/GENECONV/MaxChi/Chimaera/Bootscan/SisScan/3Seq |
| 4 | CGAa31P9 | M | 2315 | 3402 | Q32 | CGRn2616 | RDP/GENECONV/MaxChi/Chimaera/Bootscan/SisScan/3Seq |
| 5 | CGHu2 | M | 1756 | 1916 | CGRni1 | CGRn15 | RDP/GENECONV/MaxChi/Chimaera/Bootscan/SisScan/3Seq |
| 6 | CGHu2 | M | 2315 | 3402 | Q32 | CGRn2616 | RDP/GENECONV/MaxChi/Chimaera/Bootscan/SisScan/3Seq |
| 7 | CGRn93P8 | M | 2386 | 3402 | CGRn93MP8 | CGHu2 | RDP/GENECONV/MaxChi/Chimaera/Bootscan/SisScan/3Seq |
| 8 | JN131026 | M | 1413 | 2181 | JN131027 | 158577 | MaxChi/Chimaera/Bootscan/SisScan/3Seq |
| 9 | A16 | M | 2214 | 3402 | H8205 | SN7 | RDP/GENECONV/MaxChi/Chimaera/Bootscan/SisScan/3Seq |

### 3. Phylogenetic analyses and amino acid substitution analyses

The recombination isolates were excluded first. Our phylogenetic analysis showed that 11 groups were defined with S segment, each with high degree of support (Fig 2). The isolates were mainly clustered together by region. The isolates from South Korea all belonged to Group A. All the sequences from Jiangsu province and one strain from Sichuan (S85_46) province clustered in Group A with sequences from South Korea. Group B was formed with sequences collected from Russia and the Northeast of China. The isolates collected in Shaanxi were all clustered in Group C except one strain, 84FLi, which was isolated years ago in 1984. All the three isolates from Hubei constituted group D and all the isolates from Shandong clustered as a subgroup in Group E. The other subgroup of Group E contains strains collected from Anhui, Hunan, Guizhou, Shaanxi, Sichuan and Yunnan. The isolates collected from Zhejiang constitute Group G and Group H, except one strain, N8, which included in Group J. Group J also contains isolates from Jiangxi and Liaoning with the sequences of each region clustered together. Group K was formed with three isolates from Russian. We can notice that the isolates collected from Guizhou province were highly divergent. They dispersed in Group B, C, E, F and I. It indicates that HTNV has spread in Guizhou for a long period.

**Fig 2.**
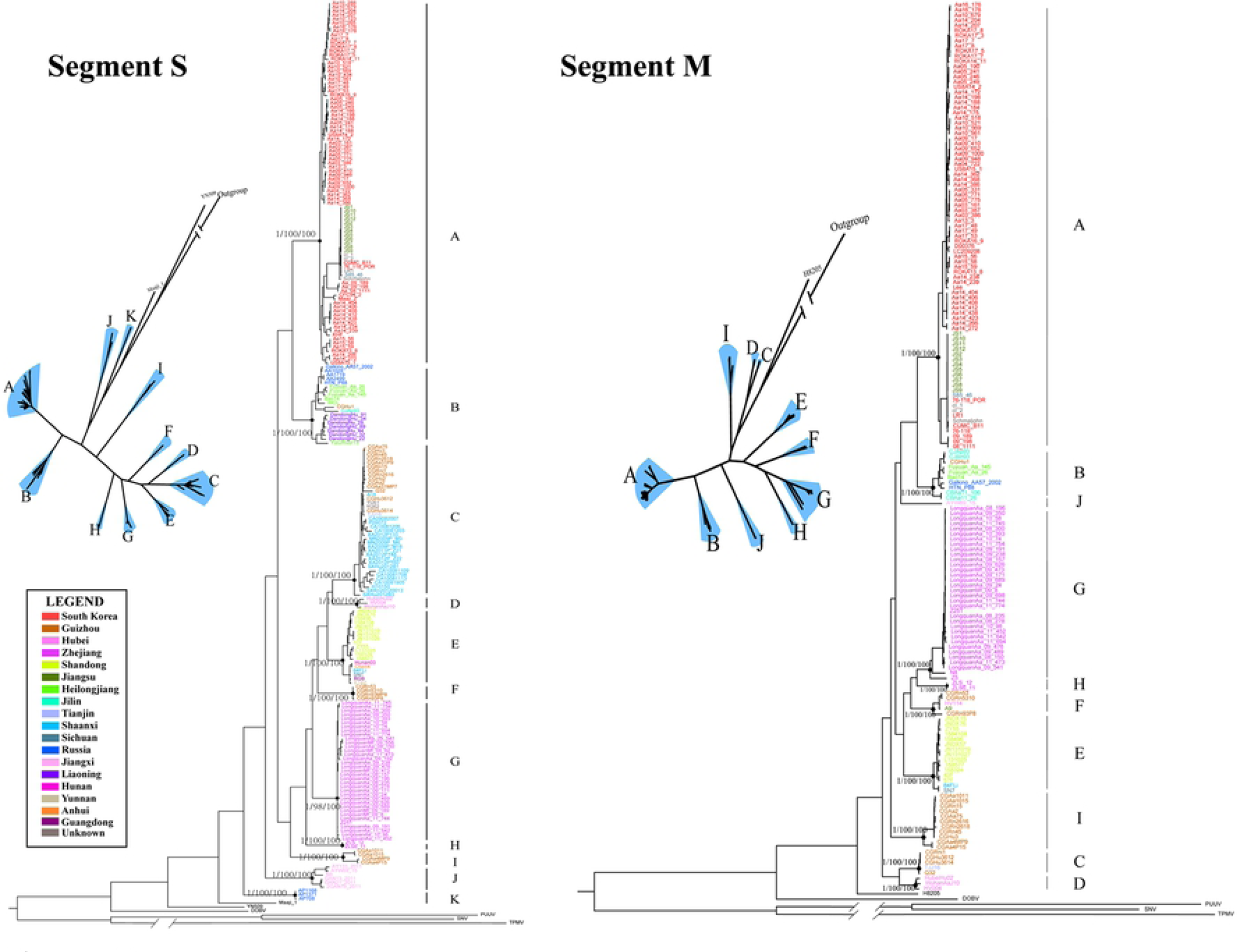
Phylogenetic trees of S gene and M gene reconstructed by MyBayes. Posterior node probabilities/SH-like approximate likelihood values/ultrafast bootstrap values for major nodes (black circles) are shown. Splits network analysis of HTNV was also shown.

There were 10 distinct phylogroups in M gene tree. The structure of phylogenetic tree reconstructed by M gene was identity with S gene tree except for 7 isolates (CGRn15, CGAa2, CGAa75, CGRn2616, CGRn2618, CGRn45, CGHu3) derived from Guizhou province. They were classified into Group I but in the S gene tree they were included in Group C. It suggested that reassortment events occurred in these strains. The branching order within the lineages differed in S and M trees, which indicates the different evolutionary history between the M and S segments.

We analyzed the variation of amino acid between the different phylogenetic groups. We noticed that 5 amino acid sites on NP gene and 37 on Gn/Gc gene were group specific and could be markers to distinguish different groups (Fig 3). No significant variant was found at the highly conserved pentapeptide motif WAASA, which is thought the cleavage of the GPC polypeptide into the Gn and Gc transmembrane proteins[29]. To determine if any of these amino acid sites were located in any known immune epitopes, we compared these significant positions against information about experimentally determined immune epitopes curated by the IEDB. We found a total of 18 positions that were located in known immune epitopes (Table 2). No significant amino acid marker was found among different hosts or collected times.

**Fig 3.**
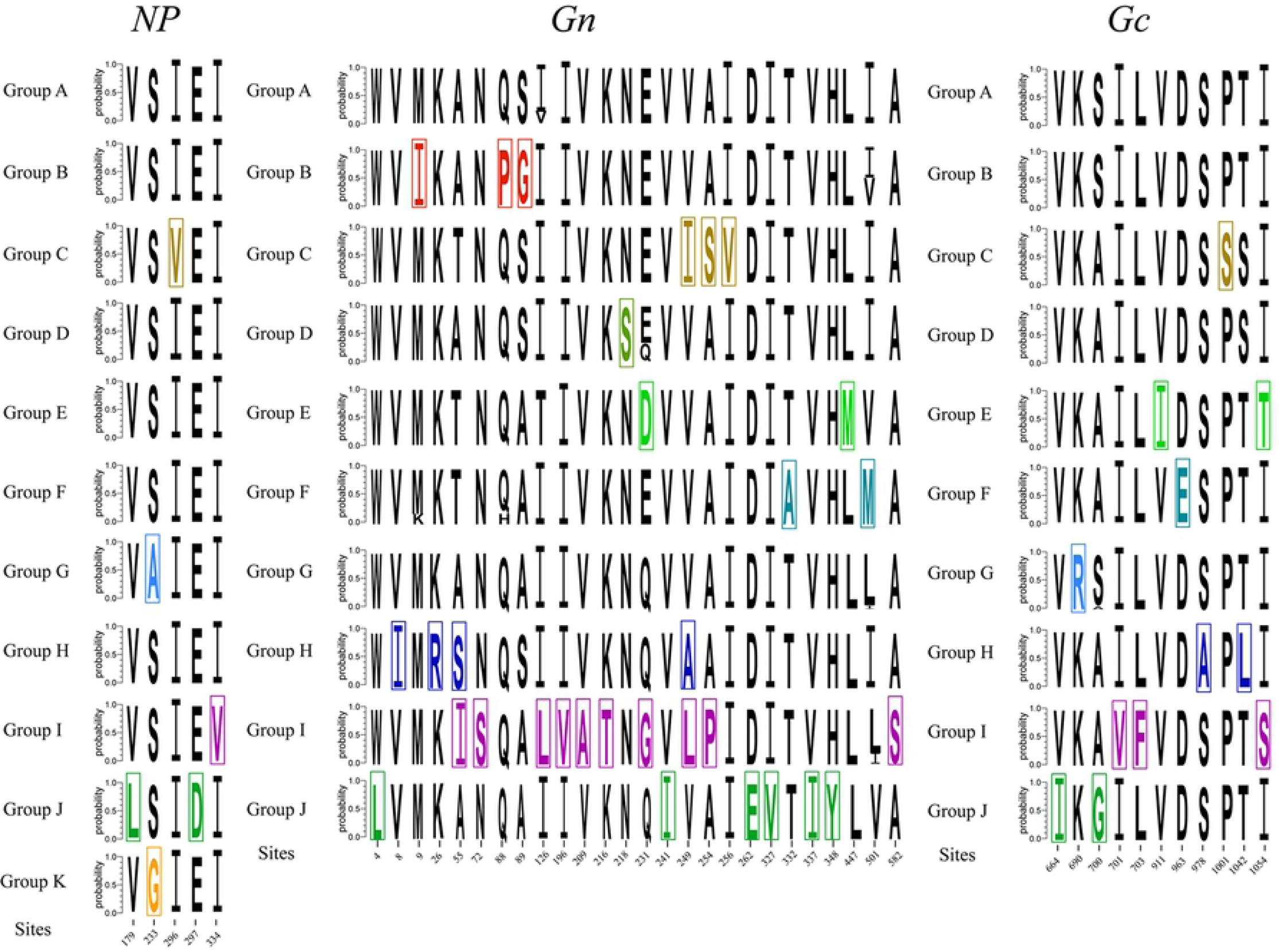
The ‘significant amino acid marker’ of each group. The specific amino acid sites of each group were marked by different colors.

**Table2.**
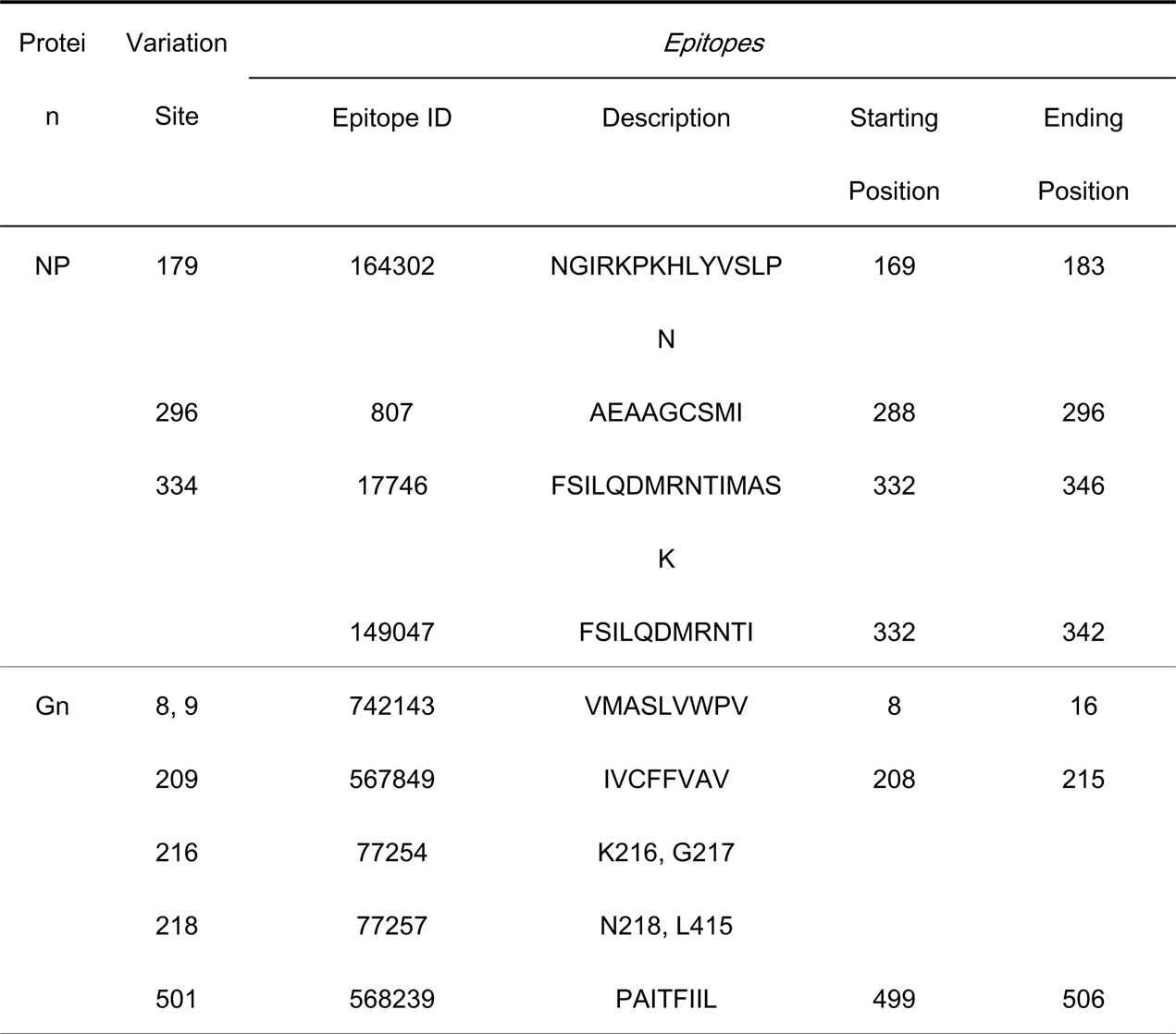
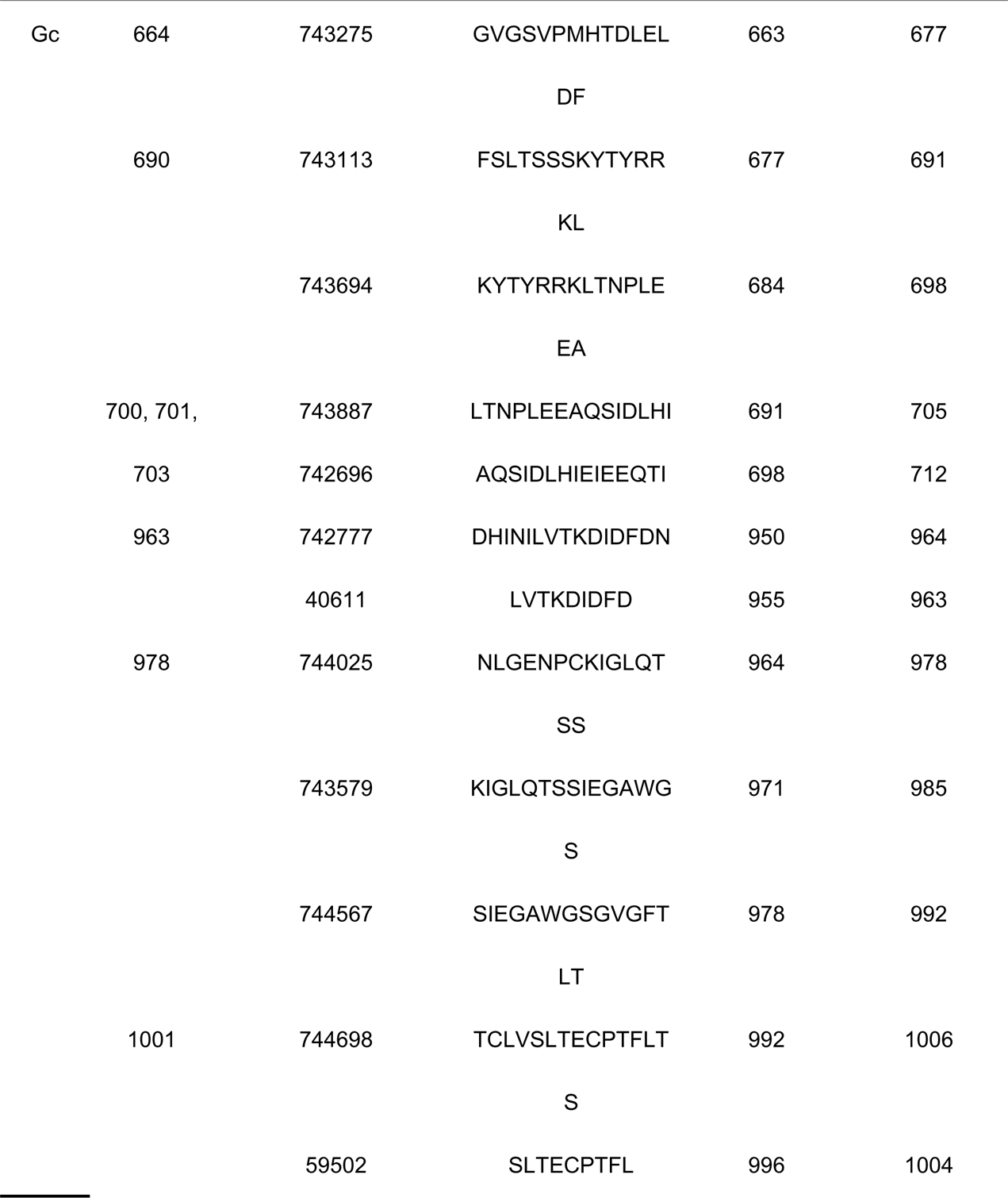
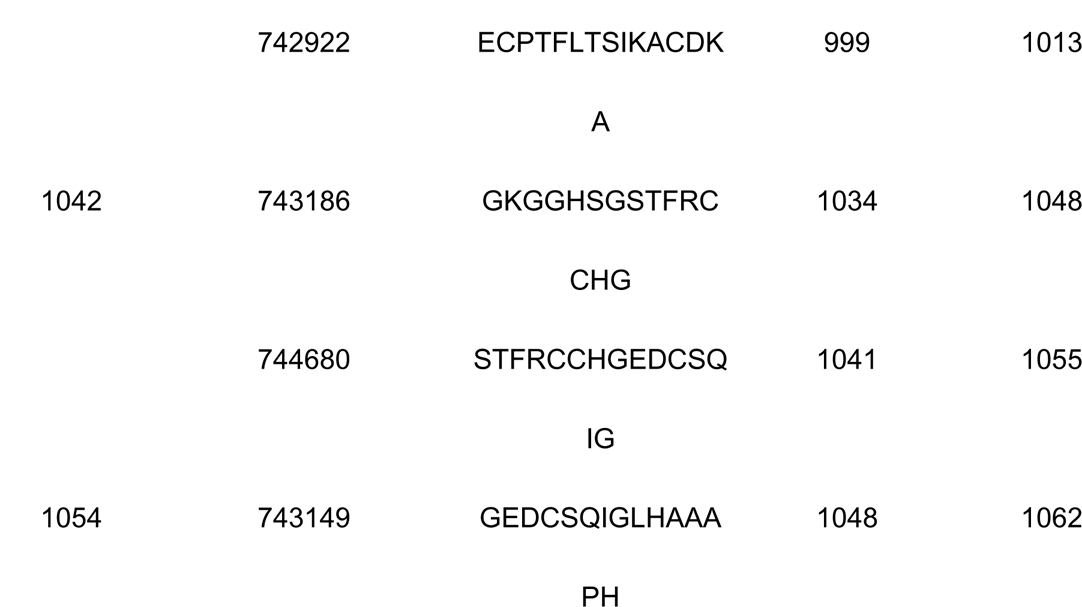
The significant sequence variations’ location on immune epitopes

| Protein | Variation Site | *Epitopes* |  |  |  |
| --- | --- | --- | --- | --- | --- |
|  |  | Epitope ID | Description | Starting Position | Ending Position |
| NP | 179 | 164302 | NGIRKPKHLYVSLP | 169 | 183 |
|  |  |  | N |  |  |
|  | 296 | 807 | AEAAGCSMI | 288 | 296 |
|  | 334 | 17746 | FSILQDMRNTIMAS | 332 | 346 |
|  |  |  | K |  |  |
|  |  | 149047 | FSILQDMRNTI | 332 | 342 |
| Gn | 8, 9 | 742143 | VMASLVWPV | 8 | 16 |
|  | 209 | 567849 | IVCFFVAV | 208 | 215 |
|  | 216 | 77254 | K216, G217 |  |  |
|  | 218 | 77257 | N218, L415 |  |  |
|  | 501 | 568239 | PAITFIIL | 499 | 506 |
| Gc | 664 | 743275 | GVGSVPMHTDLEL | 663 | 677 |
|  |  |  | DF |  |  |
|  | 690 | 743113 | FSLTSSSKYTYRR | 677 | 691 |
|  |  |  | KL |  |  |
|  |  | 743694 | KYTYRRKLTNPLE | 684 | 698 |
|  |  |  | EA |  |  |
|  | 700, 701, | 743887 | LTNPLEEAQSIDLHI | 691 | 705 |
|  | 703 | 742696 | AQSIDLHIEIEEQTI | 698 | 712 |
|  | 963 | 742777 | DHINILVTKDIDFDN | 950 | 964 |
|  |  | 40611 | LVTKDIDFD | 955 | 963 |
|  | 978 | 744025 | NLGENPCKIGLQT | 964 | 978 |
|  |  |  | SS |  |  |
|  |  | 743579 | KIGLQTSSIEGAWG | 971 | 985 |
|  |  |  | S |  |  |
|  |  | 744567 | SIEGAWGSGVGFT | 978 | 992 |
|  |  |  | LT |  |  |
|  | 1001 | 744698 | TCLVSLTECPTFLT | 992 | 1006 |
|  |  |  | S |  |  |
|  |  | 59502 | SLTECPTFL | 996 | 1004 |

|  |  |  |  |  |
| --- | --- | --- | --- | --- |
|  | 742922 | ECPTFLTSIKACDK | 999 | 1013 |
|  |  | A |  |  |
| 1042 | 743186 | GKGGHSGSTFRC | 1034 | 1048 |
|  |  | CHG |  |  |
|  | 744680 | STFRCCHGEDCSQ | 1041 | 1055 |
|  |  | IG |  |  |
| 1054 | 743149 | GEDCSQIGLHAAA | 1048 | 1062 |
|  |  | PH |  |  |

### 4. The coalescent analyses and evolution of HTNV

To avoid potential biases in the phylogeographic reconstructions due to sampling heterogeneity, we obtained a “non-redundant” subset including 51 clusters. In order to make our dataset to represent all the 15 regions (there was no sequence with exact collected year from Anhui, Sichuan and Guangdong), isolate E142, Hunan03 and JS10 were added in the dataset. Temporal evolutionary signal analyses showed a positive correlation between genetic divergence and sampling time. The strict clock with Bayesian skyline prior model yielded a higher log marginal likelihood than the others, indicating the best fit model for our dataset (Table S1).

The results of our Bayesian phylogenetic analysis showed that HTNV probably first emerged in Zhejiang province of China (root posterior probability = 0.29), with a most recent common ancestor in 1198 (95% credibility interval 593-1591; Fig 4A). The root posterior probability of Jiangxi, Shaanxi, Guizhou and Heilongjiang were also in higher levels. The MCC tree of HTNV showed four obviously two separate clusters. One was composed of isolates from the northeast of China and its surroundings, and the other was composed of isolates from the south and middle of China (Fig 4B). These indicated HTNV has been evaluating in China respectively for a long time. Reconstruction of the demographic history, using the extended Bayesian skyline plot, revealed that the HTNV population size was relatively constant from 1500 to 1970s, then expanded between 1980s and 1990s, and stayed steady from 2000 until now (Fig 3B). The mean substitution rate of 2.0427 х 10^-4^ substitution/site/year, with a 95% HPD that ranged from 1.0001 х 10^-4^ to 3.2076 х 10^-4^ was estimated.

**Fig 4.**
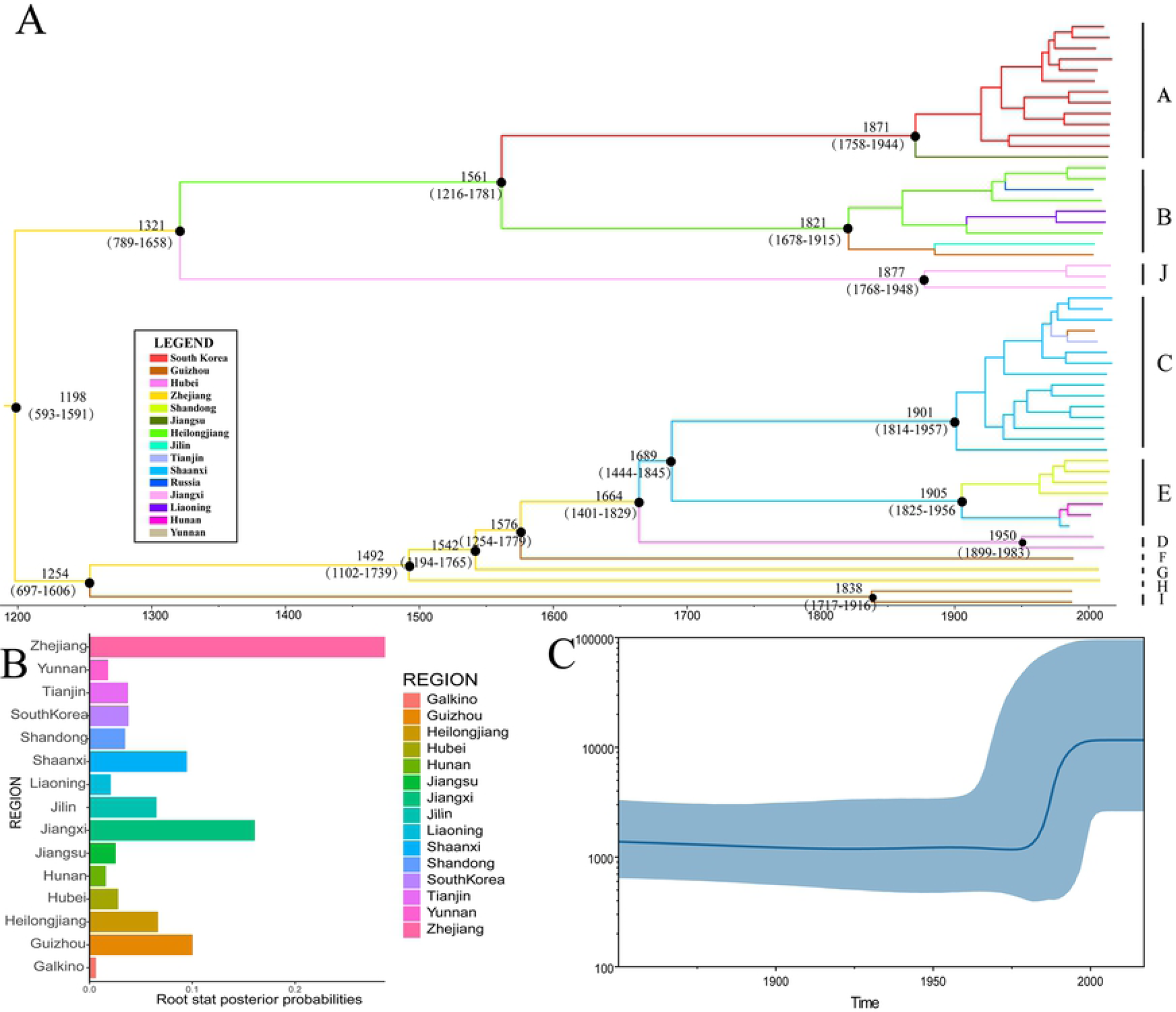
The MCC tree, root state posterior probabilities and effective population sizes of Hantaan virus. (A) MCC tree showing the evolutionary relationships and timescale of HTNV. Branch colors denote inferred location states, as shown in the legend. The most recent common ancestor and 95% credibility intervals are shown near the node. (B) The root state posterior probabilities for the different regions. (C) Bayesian skyline plot showing population size through time for HTNV. Highlighted areas correspond to 95% HPD intervals.

The spatiotemporal linkage of HTNV is shown in Fig 5. Significant location transitions were found mainly in two regions, the northeast and the middle of China (Fig 5A). It indicated that Heilongjiang may be the radiation center of the northeast regions of China and the surrounding countries. Significant migration events were found from Heilongjiang to Liaoning and Fareast of Russia. The migration from Heilongjiang to Khabarovsk was detected in high credible level (Bayes factor = 163.0). Shaanxi was assumed as the radiation center of the middle of China. Virus migration from Shaanxi to Tianjin, Yunnan and Guizhou were observed. The number of observed sate changes pinpointed Shaanxi as the main source of HTNV epidemics in China, at least in the middle of China, and Heilongjiang as the dominating source of HTNV in northeast of China (Fig 5B).

**Fig 5.**
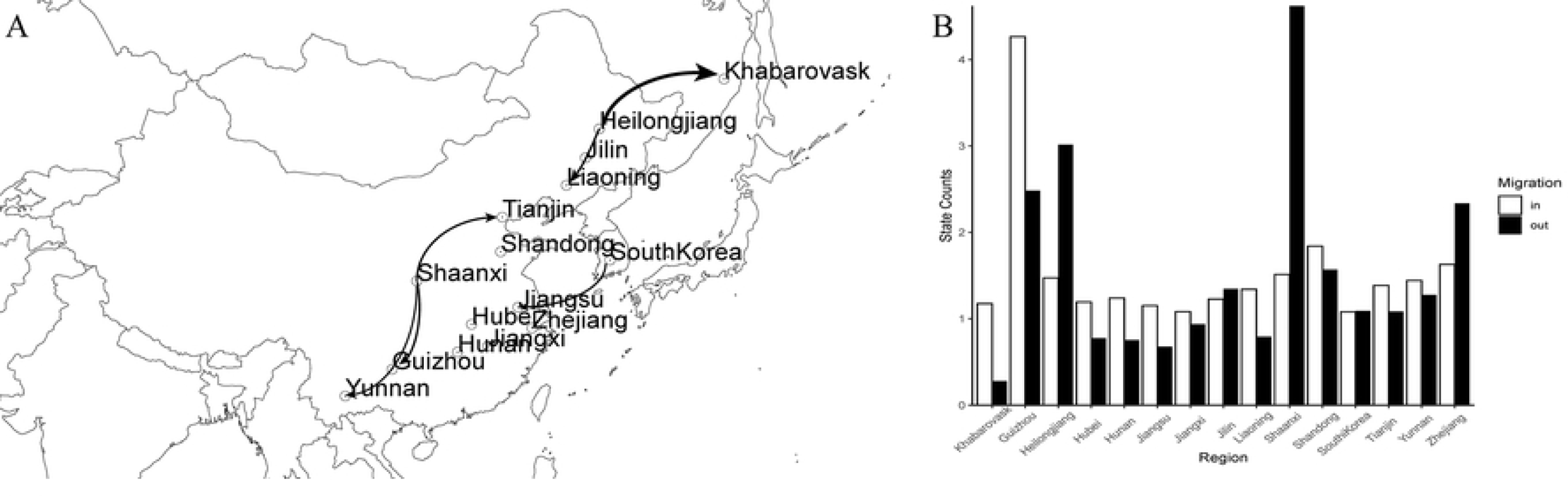
Reconstructed phylogeographic linkage and the total number of location state transitions of Hantaan virus. (A) The phylogeographic linkage constructed reconstruction of the HTNV using BEAST. Thickness of lines represents the relative transmission rate between two regions. The map was created by SpreadD3 software and the geographic data were provided by the Natural Earth (https://www.naturalearthdata.com/)*. (B) Histogram of the total number of location state transitions inferred from the HTNV dataset*.

## Discussion

In this study, we used all the whole-genome sequences of segment S and M obtained from ViPR to analyses the genetic diversity and evolution of HTNV. Our analyses show that the HTNV can be divided into 11 groups. We named the groups based on the S tree for NP gene is more stable and more sequences are available. The isolates were geographic clustering. Sequences obtained from the same geographic area clustered together. It’s worth noting that sequences from Guizhou are more diverse. We will discuss this later.

A total of 42 significant amid acid variant sites were found among the different groups. We noticed more significant variants in Group I. Isolates contained in Group I were all from Guizhou. The collection dates were during 1980s to 2000s. It also indicated a high genetic diverse of Hantaan virus in Guizhou. 18 significant amid acid variant positions were found located at known immune epitopes. Experimentation is needed to determine if peptides containing these residues can confers escape or can be used to develop serotype-specific reagents for serology-based diagnostics. None significant amino acid marker was found among different hosts and periods, demonstrating that geographic region is the mainly factor affecting the genetic diversity of HTNV. The different topologies indicate the different evolutionary history between the M and S segments, which was also found in others studies[30, 31].

Our analyses showed that both reassortment and recombination play important role in HTNV evolution. A number of studies have revealed that genetic reassortment can occur between the arthropod-borne members of the Bunyavirales naturally or experimentally[32–37]. We found 7 reassortment events occurred in M segments which is consistent with Zou, et al[8]. We found 9 recombination events in our study. All the recombination events were occurred in segment M. The reassortment and recombination events found in Gn/Gc envelope glycoproteins were consistent with their functional roles in the viral escape from immunological responses. It should be notice that recombinant A16, isolated in Shaanxi, has both major parent, H8205, and minor parent, SN7. Strain H8205 has been classified into Amur virus, which indicates recombination occurred across different species in hantavirus. Strain SN7 was isolated in Sichuan, which implies the relationship of the evolution of HTNV between Shaanxi and Sichuan. The evidences of recombination have been reported for TULV[38] and PUUV[39], even the recombination in hantavirus is rare[40].

We investigated the evolution and migration of HTNV in China and its surrounding countries using Bayesian phylogeographic inference. Our phylogenetic analyses placed the root of the tree for HTNV in Zhejiang with strong support (Fig 3B). But no migration event was found from Zhejiang to other regions. It may result from the lack of whole-genomic data from Zhejiang and its surrounding areas. Markov jump estimates between different locations pinpointed Heilongjiang as an important source of HTNV epidemics in northeast of China and far east of Russia. This finding may explain the high endemic in this area since 1931, when HFRS was first recognized in northeast of China[41, 42]. Similar pattern, Shaanxi was recognized as the origin of HTNV spread in middle of China. These can be explained by a scenario in which the virus was first introduced into these areas, then it expanded there. But the transmission sources to Heilongjiang and Shaanxi is still unclear.

Our analyses show the high diversity of isolates from Guizhou province. Phylogenetic analyses show that isolates collected from Guizhou province clustered in Group B, C, F, I. All the reassortment events and 7 of 9 recombination events occurred in Guizhou. We found same recombination pattern in strain CGAa31MP7, CGAa31P9 and CGHu2. All of these indicates that different sources of HTNV transmitted to Guizhou and evolved for a long time in this place[43, 44]. This assumption can be supported by our Markov jump estimate (Fig 4).

Guizhou is more than 1,000 meters above sea level, adding to its rich mountainous topography and as much as 92.5% of the province’s total area is characterized by mountains. It makes A. agrarius, the main host of HTNV, sympatric in rural resident areas more common. The Hengduan Mountains region was hypothesized to have played an important role in the evolutionary history of Apodemus since the Pleistocene era[45, 46].

The Yunnan-Guizhou plateau, which connects and overlaps with the Hengduan Mountains region, is also thought to play an important role in HTNV evolution. But we only found acceptable migration from Shaanxi to Guizhou in our study. This may result from the lack of sequences in Guizhou. To reduce the bias, recombinant and reassortment isolates were removed. We also got rid of similar sequences with the identity more than 99%. So limited sequences from Guizhou were used for evolutionary and phylogeographic analyses. More gene information should be collected in order to know more about the HTNV evolution and spread in this area.

The most recent common ancestor of HTNV was determined about more than 800 years ago in our study. It’s much earlier than the first HFRS patient was reported in the early 1930s[47]. But HFRS-like disease was described in Chinese writings about 1000 years ago[48], so we believe HTNV has spread in China for more than 1000 years. The most common recent ancestor by Ramsden[49] was 859 years before present (ybp) for all rodent hantaviruses estimated and 202 ybp for Murinae viruses and a mean substitution rate for rodent hantavirus of 6.67 х 10^-4^ subs/site/year with a 95% HPD that ranged from 3.86 х 10^-4^ to 9.8 х 10^-4^ subs/site/year. Partial sequences (275 nt) were used in this study, which can explain the different in tMRCA estimation. A recent study[50] showed that the estimated rate of nucleotide substitutions for the N gene of all rodent-borne hantavirus was 6.8 х 10^-4^ subs/site/year, which is similar to 6.67 х 10^-4^ subs/site/year. Our results implied HTNV evolved in a lower speed compared to the other rodent-borne hantaviruses. The estimated rate of nucleotide substitutions of DOBV and PUUV, 3 х 10^-4^ and 5.5 х 10^-4^ subs/site/year separately, supporting our hypothesis[23].

The relatively recent origin of HTNV apparently contradicts the viral-host codivergence theory. Previous estimates of evolutionary dynamics in hantaviruses were based on the critical assumption that the congruence between hantavirus and rodent phylogenies reflects codivergence between these 2 groups since the divergence of the rodent genera Mus and Rattus, approximately 10 to 40 MYA, which indicates a mean substitution rate in the range of 10^-8^ subs/site/year[39, 51, 52]. However, the observation of host–pathogen phylogenetic congruence does not necessarily indicate codivergence. Phylogenetic congruence between a parasite and its host can also arise from delayed cladogenesis, where the parasite phylogeny tracks that of the host but without temporal association[53]. This could occur if hantaviruses largely evolve host associations by cross-species transmission and related species tend to live in the same area, in which case a pattern of strong host–pathogen phylogenetic congruence could be observed in the absence of codivergence. Our evolutionary rates were estimated directly from primary sequence data sampled at known dates so that they more closely reflect the evolutionary changes undergone by the virus, at least in the short term. And with a mean substitution rate that is closer to other RNA virus, we consider our results is more reliable than codivergence theory.

Our demographic analyses revealed that HTNV population had expanded from 1980s to 1990s. As HTNV is a kind of rodent-borne virus, the population size of HTNV is relevant to rodents’ population. Previous studies have revealed that climatic factors can influence HFRS transmission through their effects on the reservoir host (mostly rodents of the family Muridae) and environmental conditions[54–56]. It’s reported that that global annual average temperatures increased by more than 1.2°F (0.7°C) from 1986 to 2016, relative to temperatures seen from 1901 to 1960[57]. Global warming may affect rodent winter survival through winter temperatures by a complicated process, and it may also influence the magnitude of HFRS outbreaks through summer climatic conditions (both temperature and rainfall). Additionally, as global climate change accelerates, the amount of annual rainfall increased accurately[58]. Moreover, agriculture improved rapidly in China at the same period, leading to more available food, which is fit for the reproduction of rodents. The steady effective population from 2000 until now is benefit from the successful strategy for HTNV prevention in China.

However, some cautions must be taken when interpreting our results. We have estimated the genetic diversity age based only on N gene sequences, the evolution rate of HTNV may be underestimated. For the Gn/Gc envelope glycoproteins are more diverse and considered evolves in a higher speed. Furthermore, there may be geographic biases due to the unbalanced of sequences collecting region. The lack of whole-genome sequences in some region may result in the lacking of knowledge about HTNV migration. In addition, lack of sequences of L gene can be obtained now. More gene information should be collected and shared to solve these problems.

In conclusion, our study revealed 11 groups of HTNV and 43 significant amino acid markers for different groups. Both recombination and reassortment events can be detected in HTNV. The origin and migration of HTNV were also shown by our analyses. And a steady effective population from 2000 until now indicated the successful strategy for HTNV prevention. Guizhou may play an important role in the evolution and spread of HTNV. As the rodents’ population and activity influence the spread and increasing of HTNV. Rodents prevention and control is an effective way to reduce the incidence of diseases. These data provide important insights for better understanding the genetic diversity and spatial-temporal dynamics of HTNV and would be useful for disease prevention and control.

---

Table S1. Log Marginal Likelihoods computed by Path Sampling and Stepping Stone Sampling

